# PTEN-loss confers dependence on the guanylate synthesis enzyme IMPDH in T-cell acute lymphoblastic leukemia

**DOI:** 10.64898/2025.12.22.696045

**Authors:** Rayees A. Padder, Thu Le Le, Emily Hyde, Maria E. D. Rubio, Eric Chiles, Namratha Sheshadri, Kelly Mulraney, Wei-Xing Zong, Xiaoyang Su, Daniel Herranz, Alexander J. Valvezan

## Abstract

Loss of the tumor suppressor PTEN is common in T-cell acute lymphoblastic leukemia (T-ALL), and is associated with poor prognosis. PTEN-loss drives robust activation of AKT/mTORC1 signaling to promote leukemic cell growth. We find that PTEN-loss in T-ALL confers dependence on the guanylate nucleotide synthesis enzyme inosine 5’-monophosphate dehydrogenase (IMPDH) for cell growth and viability. This metabolic vulnerability is dependent on sustained mTORC1 signaling and can be exploited using clinically approved IMPDH inhibitors to selectively kill PTEN-deficient T-ALL cells, and extend survival in genetic and xenograft T-ALL models in mice. Mechanistically, IMPDH inhibitors cause early DNA replication stress, followed by DNA damage. In contrast to treatment with mTORC1 inhibitors, these events culminate in robust and selective cell death in PTEN-deficient T-ALL cells. These findings reveal a targetable metabolic vulnerability in T-ALL, which could provide rationale for repurposing clinically approved IMPDH inhibitors.

**Statement of Significance:** We find that the IMPDH inhibitors mycophenolic acid and mizoribine, which are currently used as well-tolerated immunosuppressants, exert anti-leukemic activity in an aggressive molecular subset of T-ALL cells that are associated with poor prognosis. The use of clinically approved compounds to exploit this vulnerability could lead to rapid drug repurposing.

## Introduction

T-cell acute lymphoblastic leukemia (T-ALL) is an aggressive hematological malignancy that arises from the oncogenic transformation of immature T-cell progenitors. T-ALL accounts for approximately 10-15% of pediatric ALL and 25-30% of adult ALL cases (1,2). Compared with B-cell ALL, T-ALL is generally associated with higher relapse rates, and worse overall prognosis (1,2). The current standard-of-care intensive, multi-agent chemotherapy regimens yield greater than 80% overall survival in pediatric patients, but outcomes remain worse in adults, with less than 50% overall survival (3,4). Relapse is associated with very poor prognosis, with a 5-year survival rate less than 25% (5–9).

The tumor suppressor PTEN is inactivated or downregulated in at least 15-25% of T-ALL, which is associated with higher relapse rates and reduced overall survival in children and adults (10–18). PTEN is a lipid phosphatase that dephosphorylates PIP3 to PIP2, antagonizing the activity of PI3K. PTEN-loss causes accumulation of PIP3, leading to activation of AKT. AKT then phosphorylates many factors to influence metabolism, growth, proliferation, and survival (19,20). A critical effector downstream of AKT is the mechanistic Target of Rapamycin Complex 1 (mTORC1), which AKT activates by inhibiting an essential negative regulator of mTORC1 called the TSC complex (19,20). mTORC1 is a master metabolic regulator that stimulates synthesis of the major macromolecules required for cell growth and proliferation, including proteins, lipids, and nucleic acids (21–23). Underscoring its central importance in coordinating anabolic growth, mTORC1 is activated in the majority of human cancers across nearly all lineages (24). This occurs downstream of many of the most commonly mutated oncogenes and tumor suppressors in human cancers, including receptor tyrosine kinases, the PI3K/Akt pathway, and the Ras pathway, which converge to activate mTORC1 by inhibiting the TSC complex (25). There has been intense interest in targeting mTORC1 in cancer, but the mTORC1 inhibitors rapamycin and its analogs have shown limited efficacy as single agent therapies, generally inducing cytostatic, rather than cytotoxic, responses (26,27).

As an alternative strategy to targeting mTORC1 itself in cancer, we discovered a metabolic vulnerability induced downstream of mTORC1 that allows selective killing of tumor cells that have uncontrolled mTORC1 activation (28,29). In tumors with uncontrolled mTORC1 activation caused by loss of the TSC complex, we found that mTORC1 strongly increases the cellular demand for nucleotides by driving the synthesis of ribosomal RNA (rRNA) for ribosome biogenesis. This confers dependence on *de novo* nucleotide synthesis pathways, which are also stimulated by mTORC1 in parallel (30–32), to sustain sufficient free NTP pools. This unique metabolic dependency can be exploited using clinically approved inhibitors of the rate-limiting enzyme in *de novo* guanylate nucleotide synthesis, inosine 5’-monophosphate dehydrogenase (IMPDH), which have been used for decades as safe and well-tolerated immunosuppressants (28,29,33,34). IMPDH inhibitors demonstrate robust efficacy in genetic and xenograft models of TSC complex-deficient tumor growth (28,29). IMPDH has also emerged as a potential target for anti-cancer therapy in preclinical models of small-cell lung cancer (35,36), hepatocellular carcinoma (37–39), and glioblastoma (40–42), with early-stage clinical trials in glioblastoma recruiting (NCT05236036) and underway (NCT04477200).

Our previous studies targeting IMPDH in TSC complex-deficient cells and tumors used two clinically approved IMPDH inhibitors: mizoribine and mycophenolic acid (MPA). Mizoribine (Bredinin) is used clinically in Asia as an immunosuppressant for organ transplantation, as well as for the treatment of autoimmune disorders. Mycophenolic acid is used in the US and Europe for the same indications, and is commonly administered as an FDA-approved orally bioavailable pro-drug mycophenolate mofetil (MMF/CellCept) (34). Mizoribine is superior to MPA/MMF in targeting TSC complex-deficient tumors *in vivo* (29). However, mizoribine is a nucleoside analog that can only inhibit IMPDH when phosphorylated to generate the nucleotide analog mizoribine monophosphate (43,44). This is carried out by adenosine kinase (ADK), and cells with low ADK-expression are relatively insensitive to mizoribine (28,29). ADK levels do not affect sensitivity to MPA/MMF (29). Thus, each IMPDH inhibitor has advantages and disadvantages in different settings.

Given the frequency of mTORC1 activation in cancer, we sought to determine whether mTORC1-activating mutations that lie upstream of the TSC complex could confer robust dependence on IMPDH. Here we find that PTEN-loss sensitizes T-ALL cells to IMPDH inhibition, in an mTORC1-dependent manner, and that clinically approved IMPDH inhibitors extend survival in genetic and xenograft T-ALL models in mice. Our data also suggest that other mTORC1-activating mutations could similarly confer strong sensitivity to IMPDH inhibition in T-ALL. Together, these findings establish a targetable metabolic vulnerability in T-ALL, and significantly broaden the potential anti-cancer applications for IMPDH inhibitors.

## Results

### IMPDH inhibitors induce cell death in *PTEN*-deficient T-cell acute lymphoblastic leukemia cells

To identify the cancer types with the highest dependence on IMPDH for growth and viability, we examined The Cancer Dependency Map CRISPR screening data (DepMap, https://depmap.org/portal/). Consistent with the known immunosuppressive effects of IMPDH inhibitors, leukemias and lymphomas were most dependent on IMPDH2, with T-cell acute lymphoblastic leukemia scoring at the top of the list (Supplemental Figure S1A). Further stratification of cell lines based on *PTEN*-status revealed that, despite strong overall dependence on IMPDH2, *PTEN*-mutant cell lines were generally more dependent compared to *PTEN*-wild type (Supplemental Figure S1B). To determine the response to pharmacological inhibition of IMPDH, a panel of nine human T-ALL cell lines were treated for 3 days with a dose curve of the FDA-approved IMPDH inhibitor mycophenolic acid (MPA). PTEN-deficient cells were generally more sensitive to MPA compared to PTEN-expressing cells (Figure 1A, B). Similar results were observed using the IMPDH inhibitor mizoribine, as well as the recently discovered IMPDH inhibitor gliocidin (42) (Supplemental Figure S1C, S1D). Immunoblot analysis revealed robust, serum/growth factor-independent activation of Akt/mTORC1 signaling in the PTEN-deficient cells, as indicated by activating phosphorylation of Akt (S473 and T308) and phosphorylation of the direct Akt substrate GSK-3 (S9/21), as well as phosphorylation of the direct mTORC1 substrates S6 Kinase (T389) and 4E-BP1 (multi-site phosphorylation resulting in upward mobility shifts (Figure 1C,D).

**Figure 1:**
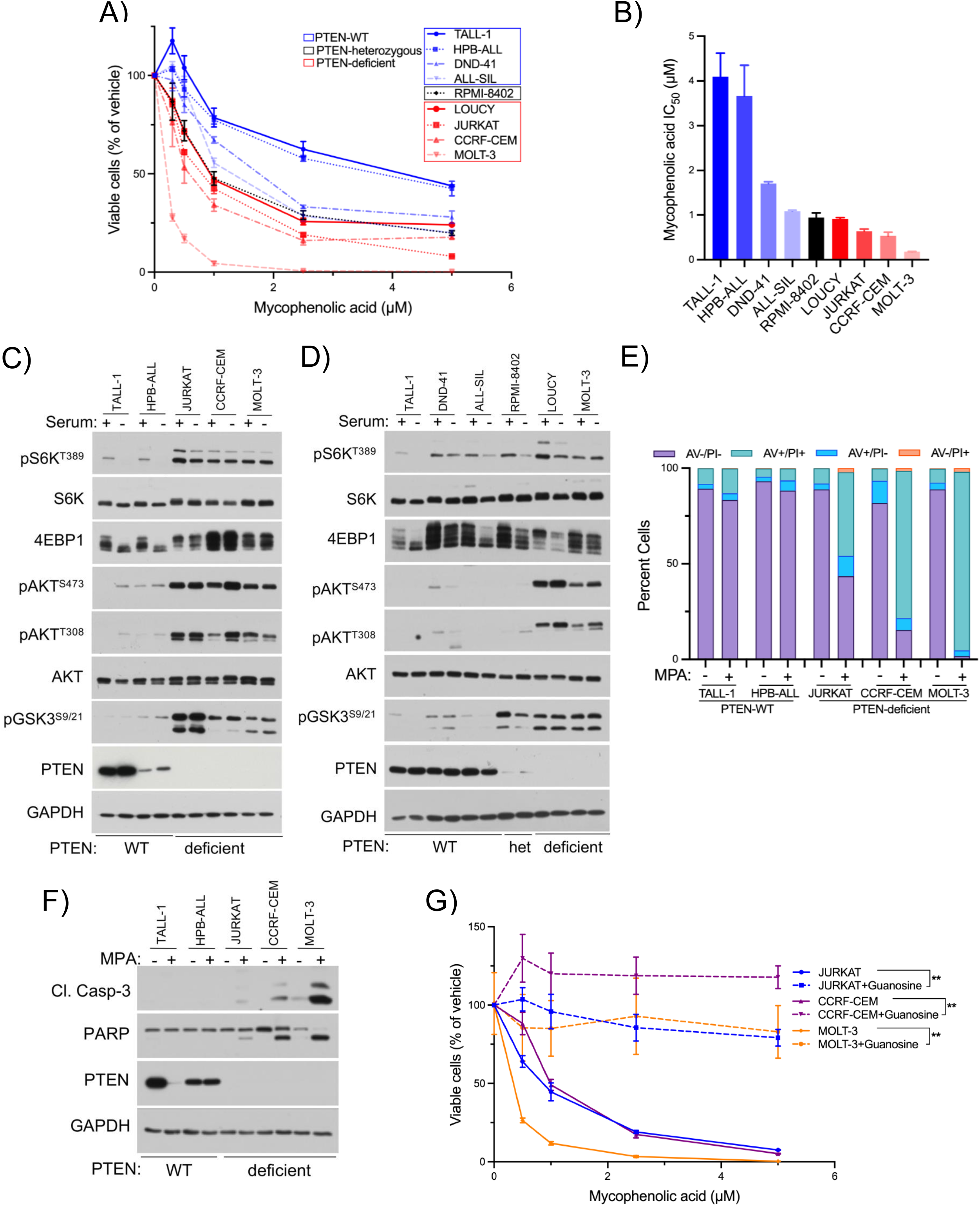
IMPDH inhibitors induce cell death in *PTEN*-deficient T-cell acute lymphoblastic leukemia cells. (**A, B)** Indicated human T-ALL cells were treated for 72 hrs with indicated concentrations of the FDA approved IMPDH inhibitor mycophenolic acid. **(A)** Viable cells were measured using the Cell Titer Glo Cell Viability Assay (Promega), graphed as percent of vehicle-treated cells. **(B)** Mycophenolic acid IC_50_ values, calculated from the experiment in **(A).** **(C, D)** T-ALL cells from **(A)** grown in complete media or serum starved for 18 hours followed by immunoblotting for indicated proteins. **(E)** Indicated T-ALL cells treated for 72 hrs with vehicle or mycophenolic acid (MPA, 2.5 µM). Cell death was quantified by Annexin V/ Propidium Iodide (AV/PI) staining and graphed as percent of the total cell population. **(F)** Immunoblots from indicated T-ALL cells treated with vehicle or MPA (1 µM, 48 hrs). **(G)** Indicated cells treated with indicated doses of MPA, with or without exogenous guanosine (25 µM). Viable cells were measured as in **(A)** and graphed as percent of vehicle-treated cells. Graphical data are represented as mean ± SEM. **p < 0.0001 by two-way ANOVA. (See also Figure S1).

To quantify the cell-killing effects of IMPDH inhibition, PTEN-wild type T-ALL1 and HPB-ALL cells that have intact, serum/growth factor-dependent mTORC1 signaling, and PTEN-deficient JURKAT, CCRF-CEM and MOLT-3 cells that have uncontrolled, serum/growth factor-independent mTORC1 activation (Figure 1C), were treated with MPA followed by Annexin V/propidium iodide staining (AV/PI). MPA induces robust and selective cell death in the PTEN-deficient cells, including over 98% cell death in MOLT3 cells, with minimal effects on the PTEN-wild type cells (Figure 1E). This is accompanied by Caspase-3 cleavage and cleavage of the Caspase-3 substrate PARP (Figure 1F). The cell-killing effects of MPA are rescued by co-treatment with the pan-Caspase inhibitor Q-VD-OPh (Supplemental Figure S1E), indicating that cell death is occurring primarily through apoptosis. Supplementation with exogenous guanosine, which can be converted to guanylate nucleotides through the purine salvage pathway independent of IMPDH (45), rescues cell proliferation, viability and apoptosis, confirming that the effects of MPA are through guanylate depletion (Figure 1G, Supplemental Figure S1F). Thus, IMPDH inhibition induces robust and selective cell death in PTEN-deficient T-ALL cells.

### PTEN-deficiency confers sensitivity to IMPDH inhibitors through active mTORC1 signaling

To determine whether PTEN-loss plays a causal role in sensitizing T-ALL cells to IMPDH inhibition, we used JURKAT cells engineered with a doxycycline-inducible PTEN (46). Doxycycline treatment induced PTEN expression, reduced Akt/mTORC1 signaling (Supplemental Figure S2A), and correspondingly reduced cleavage of Caspase-3 and PARP in response to IMPDH inhibition (Figure 2A). Conversely, we deleted PTEN in PTEN-expressing Ba/F3 cells via CRISPR-Cas9, which increased sensitivity to IMPDH inhibition when measuring viable cells, induction of apoptosis, and cell death (Figure 2B-D). Thus, PTEN-deficiency confers sensitivity to IMPDH inhibition.

**Figure 2:**
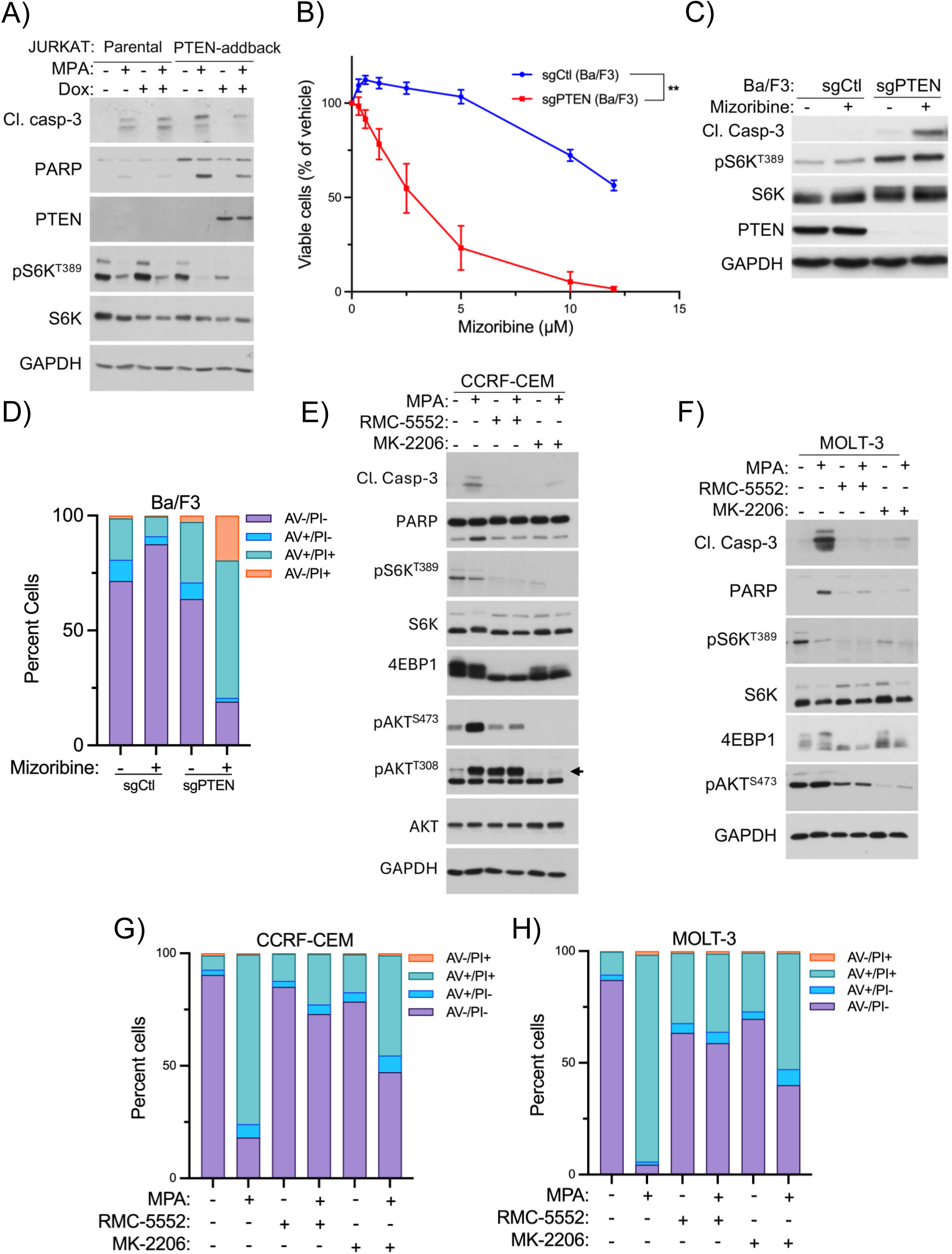
PTEN-deficiency confers sensitivity to IMPDH inhibitors through active mTORC1 signaling. **(A)** Immunoblots from parental Jurkat cells and Jurkat cells with doxycycline-inducible PTEN-addback, treated with mycophenolic acid (MPA, 1 µM, 48 hrs) in the presence or absence of doxycycline (Dox, 1 µg/mL), following 24 hr doxycycline pre-treatment. **(B-D)** Ba/F3 cells with CRISPR control (sgCtl) or *Pten* deletion (sgPTEN) were **(B)** treated with indicated doses of the IMPDH inhibitor mizoribine for 72 hours. Viable cells were measured using the Cell Titer Glo Cell Viability Assay and graphed as percent of vehicle-treated cells. **(C)** Immunoblots on indicated proteins treated with vehicle or mizoribine (5 µM, 48 hrs). **(D)** Ba/F3 cells treated with vehicle or mizoribine (5 µM, 72 hrs). Cell death was quantified by Annexin V/ Propidium Iodide (AV/PI) staining and graphed as percent of the total cell population. **(E, F)** Immunoblots from the indicated PTEN-deficient T-ALL cells treated for 48 hrs with MPA (1 µM) with or without the mTORC1 inhibitor RMC-5552 (1 nM) or the AKT inhibitor MK-2206 (2 µM). **(G, H)** Indicated cells treated for 72 hrs with MPA (2.5 µM) with or without RMC-5552 (1 nM) or MK-2206 (2 µM). Cell death was quantified and graphed as in **(D)**. Graphical data are represented as mean ± SEM. **p < 0.0001 by two-way ANOVA. (See also Figure S2).

To determine the role of Akt/mTORC1 signaling in the response to IMPDH inhibition, PTEN-deficient CCRF-CEM and MOLT-3 cells were treated with MPA with or without the bi-steric mTORC1 inhibitor RMC-5552 (47) or the AKT inhibitor MK-2206. Inhibition of mTORC1 or Akt blocked MPA-induced apoptosis, measured by cleaved caspase-3 and cleaved PARP (Figures 2E, 2F), and cell death measured by Annexin V/PI staining (Figures 2G, 2H). This effect was more pronounced with RMC-5552 compared to MK-2206, corresponding with greater mTORC1 inhibition upon RMC-5552 treatment (Figure 2E, 2F evident by 4E-BP1 mobility shift). In JURKAT cells, mTORC1 inhibition reduced MPA-induced apoptosis, whereas Akt inhibition did not (Supplemental Figure S2B, S2C), suggesting other factors in these cells may contribute to IMPDH inhibitor sensitivity in an mTORC1-dependent manner. Together, these data indicate that mTORC1 activation in PTEN-deficient T-ALL cells promotes sensitivity to IMPDH inhibition.

### IMPDH inhibition causes DNA replication stress and DNA damage in PTEN-deficient T-ALL cells

To determine the molecular events preceding cell death, PTEN-deficient JURKAT, CCRF-CEM and MOLT-3 cells were treated with a time course of MPA. In all 3 cell lines, MPA induced phosphorylation of the cell cycle checkpoint protein CHK1 (S345), as well as RPA (T21), markers of DNA replication stress (48) (Figure 3A). This effect was rescued by supplementation with exogenous guanosine, indicating that it stems from guanylate depletion upon MPA treatment (Supplemental Figure S3A). MPA also increased Cyclin E levels, suggesting accumulation of cells in S-phase of the cell cycle (Figure 3A), which was confirmed by flow cytometry for propidium iodide staining (Figure 3B). These effects are consistent with induction of the intra-S-phase checkpoint, a known response to DNA replication stress that results in phosphorylation and activation of CHK1 which arrests cells in S-phase (48). MPA-induced DNA replication stress was blocked by co-treatment with RMC-5552, and to a lesser extent MK-2206, which was correspondingly less effective at inhibiting mTORC1, as above (Figure 3D, 3E).

**Figure 3:**
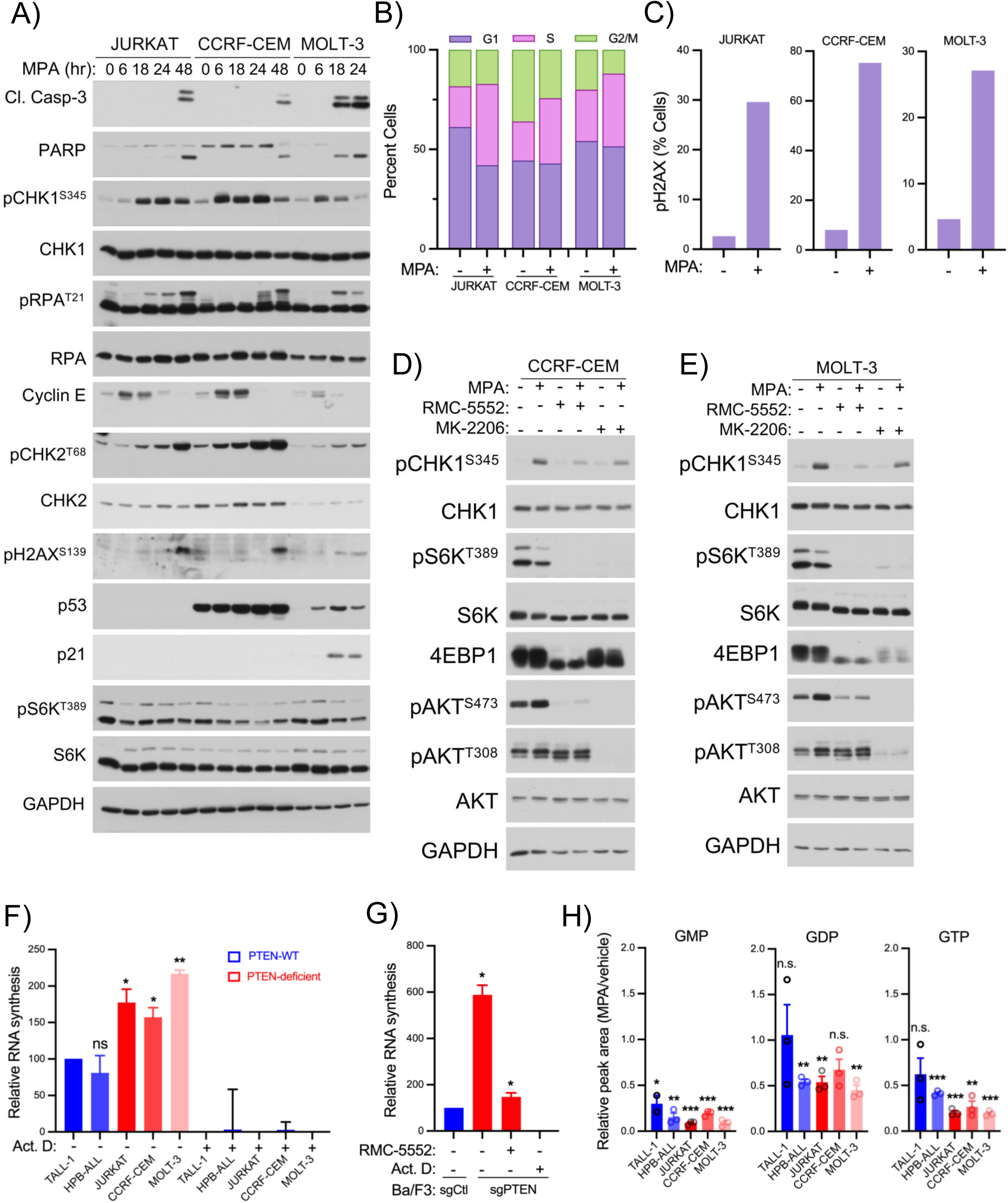
IMPDH inhibition causes DNA replication stress and DNA damage in PTEN-deficient T-ALL cells. **(A)** Immunoblots from indicated cells treated with mycophenolic acid (MPA, 1 µM) for indicated times. **(B)** Indicated cells were treated with MPA (1 µM, 24 hrs). Cell cycle distribution was quantified by flow cytometry for propidium iodide staining. **(C)** Indicated cells were treated with MPA (1 µM, 16 hrs) followed by flow cytometry for phosho-H2AX^S139^ immunostaining, graphed as percent of positive cells. **(D, E)** Immunoblots on indicated cells treated for 24 hrs with MPA (1 µM) with or without RMC-5552 (1 nM) or MK-2206 (2 µM). **(F, G)** RNA synthesis measured in indicated cells using the RNA Synthesis Assay kit (Abcam). Cells were treated for 3 hrs with vehicle or actinomycin D (Act. D) and then labeled for 1 hr with 5-ethynyl-uridine (EU), followed by quantification by flow cytometry. **(H)** Relative abundance of the indicated metabolites, measured by LC-MS/MS, in cells treated with MPA (1 µM, 6 hours), and graphed normalized to vehicle-treated cells. Graphical data are represented as mean ± SEM. *p < 0.05, **p < 0.005, ***p < 0.0005 by unpaired Welch’s t-test. (See also Figure S3).

Prolonged, unresolved DNA replication stress can result in DNA damage (48). Induction of the DNA damage marker γH2AX (phospho-histone H2AX S139) was observed at later time points correlating with the induction of apoptosis (Figure 3A, 3C). Activating phosphorylation of CHK2 (T68), which occurs in response to DNA damage, was also observed (Figure 3A). Together these data indicate that IMPDH inhibition causes DNA replication stress and DNA damage in PTEN-deficient T-ALL cells.

In MOLT-3 cells, but not JURKAT or CCRF-CEM cells, MPA increased the levels of p53 and its transcriptional target p21 (Figure 3A). Compared to JURKAT and CCRF-CEM cells, MOLT-3 cells exhibit greater sensitivity to IMPDH inhibitors (Figure 1A, Supplemental Figures S1C, S1D), and earlier induction of apoptosis (Figure 3A), thus correlating with the presence of functional p53 in these cells.

Ribosomal RNA (rRNA) comprises 80-85% of total cellular RNA in most mammalian cell types, and mTORC1 drives ribosomal RNA (rRNA) synthesis for ribosome biogenesis (49–51). Previous work demonstrated that increased RNA synthesis rates can enhance cellular demand for nucleotides, thus conferring dependence on IMPDH to meet the elevated demand (28,29,36,40). We measured RNA synthesis rates via 5-ethynyl-uridine (5-EU) labeling. RNA synthesis rates in PTEN-deficient cells were 60%-125% greater compared to PTEN-expressing cells (Figure 3F). PTEN-deficient Ba/F3 cells had even more strongly elevated rates of RNA synthesis compared to isogenic control cells, and this difference was abolished by RMC-5552 treatment, demonstrating that it is driven by mTORC1 (Figure 3G). Consistent with increased GTP utilization, 6 hr treatment with MPA reduced GTP levels to a greater extent in PTEN-deficient compared to PTEN-expressing T-ALL cells, measured by LC-MS/MS-based metabolite profiling (Figure 3H). This difference was not clearly evident with GMP or GDP levels, which were depleted similarly by MPA across cell lines, possibly because they can be rapidly converted to GTP in an attempt to maintain GTP pools (Figure 3H). MPA did not strongly affect ATP levels, but caused accumulation of the IMPDH substrate IMP, as well as intermediates upstream of IMP in the *de novo* purine synthesis pathway (AICAR) and salvage pathway (inosine, hypoxanthine) (Supplemental Figures S3B – S3G). Despite approximately 20-60-fold accumulation of hypoxanthine, MPA did not cause accumulation of xanthine (Supplemental Figure S3G, S3H). Together, these data suggest that in PTEN-deficient T-ALL cells, increased mTORC1-driven RNA synthesis increases dependence on IMPDH to sustain GTP pools, and that IMPDH inhibition causes early DNA replication stress and later DNA damage, culminating in apoptosis.

### Clinically approved IMPDH inhibitors extend survival in genetic and xenograft models of PTEN-deficient T-ALL

We next asked whether IMPDH inhibition could be effective against PTEN-deficient T-ALL *in vivo*, using an established genetically engineered mouse model (52). To induce T-ALL development, bone marrow hematopoietic progenitors from tamoxifen-inducible conditional *Pten* knockout mice (*Rosa26*^Cre-ERT2/+^*Pten*^fl/fl^) were infected with retrovirus expressing a constitutively active oncogenic NOTCH1 receptor mutant (L1601P Δ-PEST) along with a luciferase reporter, and then transplanted into isogenic recipient mice. Activating mutations in the Notch signaling receptor NOTCH1 are present in greater than 60% of human T-ALL and frequently co-occur with PTEN-loss (10,16,18,53). T-ALL cells from these mice were then injected into secondary recipients (Day 0) and treated with tamoxifen 2 days later (Day 2) to generate *Pten*-deleted T-ALL, as previously described (52). 4 days after tamoxifen injection (Day 6), we began daily treatment with vehicle or mizoribine (100 mg/kg/day for the first week and then 75 mg/kg/day thereafter). In the vehicle group, mice rapidly lost weight as the T-ALL progressed, and all mice succumbed between Day 11-14 (Figure 4A, Supplemental Figure S4A). Despite starting treatment on Day 6, when vehicle mice were only 5 days from succumbing, mizoribine treatment significantly extended survival and delayed T-ALL-induced weight loss (Figure 4A, Supplemental Figure S4A). Luciferase imaging on Day 11 showed that mizoribine reduced T-ALL burden by 60%, and prevented the T-ALL cell accumulation in the thymus that occurs in vehicle-treated mice (Figure 4B, 4C). In mice with *Pten*-expressing, NOTCH1-driven T-ALL (no tamoxifen injection on Day 2), mizoribine also extended lifespan, though the effect was less pronounced compared to mice with *Pten*-deleted T-ALL (Supplemental Figure S4B, S4C). This is consistent with the moderate efficacy of IMPDH inhibition and IMPDH2 dependence in some PTEN-expressing T-ALL cell lines (Figure 1A, 1B, Supplemental Figure S1B). We also tested the FDA-approved orally bioavailable prodrug of MPA, mycophenolate mofetil (MMF, CellCept). However, in line with our previous report of superior efficacy of mizoribine in some tumor models (29), MMF did not robustly extend lifespan in mice bearing *Pten*-deleted or *Pten*-expressing T-ALL (Supplemental Figure S4B, S4D).

**Figure 4:**
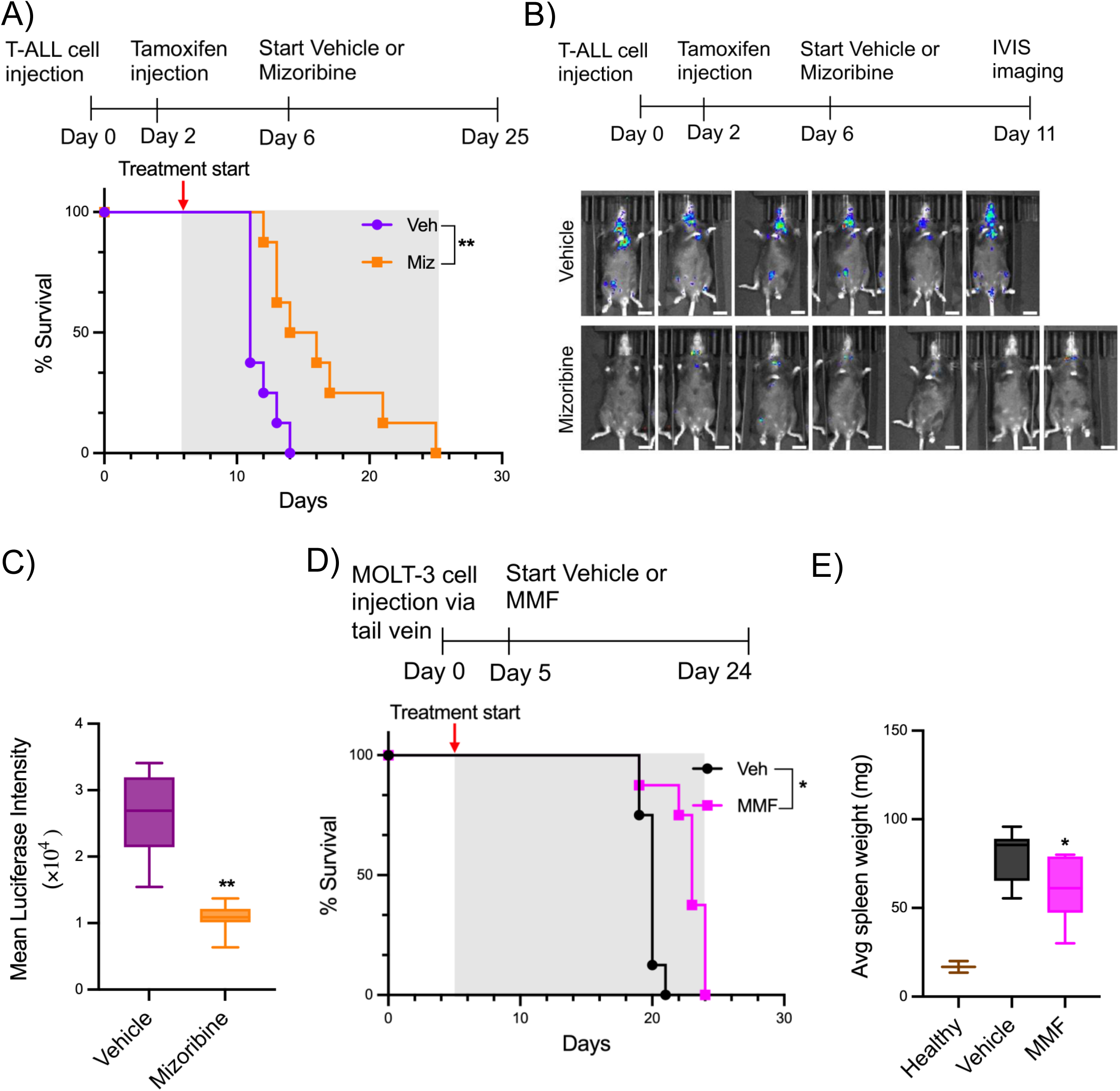
Clinically approved IMPDH inhibitors extend survival in genetic and xenograft models of PTEN-deficient T-ALL. **(A-C)** To induce T-ALL development, bone marrow hematopoietic progenitors from tamoxifen-inducible conditional PTEN-knockout mice (Rosa26^Cre-ERT2/+^PTEN^fl/fl^) were infected with retrovirus expressing a constitutively active oncogenic NOTCH1 receptor mutant (L1601P Δ-PEST), along with a luciferase reporter, and then transplanted into isogenic recipient mice. T-ALL cells from these mice were then injected into irradiated secondary recipients (via retro-orbital injection) and treated with tamoxifen 2 days later to generate PTEN-deleted T-ALL. Beginning 4 days after tamoxifen injection, mice were treated daily with vehicle or mizoribine (100 mg/kg/day for the first week and then 75 mg/kg/day thereafter) until succumbing. **(A)** Kaplan-Meier survival plots. n=8 mice per group. **(B)** IVIS imaging after 5 days of treatment with vehicle or mizoribine. **(C)** Quantification of mean luciferase intensity in **(B)**. **(D, E)** T-ALL xenografts were established via tail vein injection of MOLT-3 cells. Beginning 5 days later, mice were treated daily with vehicle or mycophenolate mofetil (MMF, 200 mg/kg/day) until succumbing. **(D)** Kaplan-Meier survival plots. n=8 mice per group. **(E)** Average spleen weights from vehicle and MMF-treated mice compared to healthy non-leukemic controls. *p < 0.05, **p < 0.005 by unpaired Welch’s t-test or Mantel-Cox test. (See also Figure S4).

In a xenograft T-ALL model established via tail vein injection of MOLT-3 cells, which are strongly sensitive to MPA in culture (Figure 1A, 1B, 1E), MMF modestly but significantly extended survival, and partially rescued the splenomegaly, but not hepatomegaly, that occurs in this model (Figure 4D, 4E, Supplemental Figure S4E, S4F). MOLT-3 cells express low levels of ADK, which is required for mizoribine efficacy (Supplemental Figure S4G), and thus were insensitive to mizoribine *in vivo* (Supplemental Figure S4H). Together, these data indicate that IMPDH inhibition can reduce T-ALL burden and extend lifespan *in vivo,* but that efficacy is strongly influenced by the choice of IMPDH inhibitor, with mizoribine and MPA/MMF having advantages and disadvantages in different models. MMF can show modest efficacy, which is superior to mizoribine when ADK levels are low, but mizoribine can strongly outperform MMF when ADK levels are not limiting.

### Uncontrolled mTORC1 activation in T-ALL cells with wild-type PTEN confers sensitivity to IMPDH inhibition

Given our observation that mTORC1 activation downstream of PTEN-deficiency confers sensitivity to IMPDH inhibition, we asked whether this susceptibility was strictly restricted to PTEN-loss. We identified two PTEN-wild type T-ALL cell lines, HSB-2 and KOPTK-1, which exhibited similar sensitivity to MPA as PTEN-deficient cells (Figure 5A). Despite robust PTEN expression, mTORC1 signaling in these cells was serum/growth factor-independent (Figure 5B), indicating that they have other mTORC1-activating mutations. MPA strongly induces cell death in these cells, which is rescued by the mTORC1 inhibitor RMC-5552 (Figure 5C). As in PTEN-deficient cells, MPA also induces DNA replication stress, S-phase delay/arrest, DNA damage, and apoptosis (Figure 5D, 5E). Similar to MOLT-3 cells, p53 was induced in KOPTK-1 cells (but not HSB-2 cells), correlating with earlier induction of apoptosis (Figure 5E). MPA caused guanylate nucleotide depletion without strongly affecting ATP levels, and caused accumulation of IMP, inosine and hypoxanthine, but not xanthine (Supplemental Figure S5A-F).

**Figure 5:**
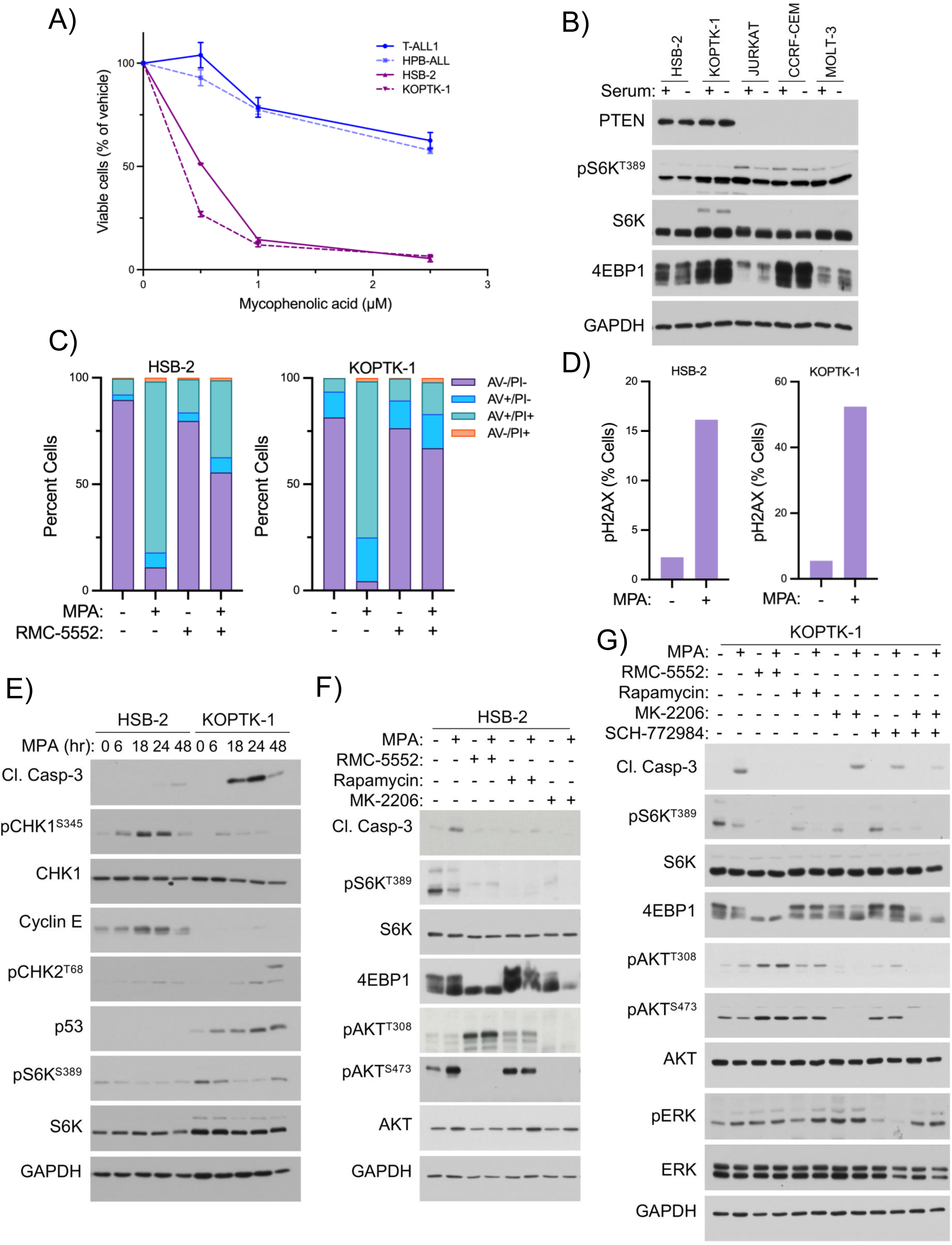
Uncontrolled mTORC1 activation in T-ALL cells with wild-type PTEN confers sensitivity to IMPDH inhibition. **(A)** Indicated cells were treated for 72 hrs with indicated concentrations of mycophenolic acid. Viable cells were measured using the Cell Titer Glo Cell Viability Assay (Promega) and are graphed as percent of vehicle-treated cells. **(B)** Immunoblots of indicated cells grown in complete media or serum starved for 18 hours. **(C)** Indicated cells treated for 72 hrs with MPA (2.5 µM) with or without RMC-5552 (1 nM). Cell death was quantified by Annexin V/ Propidium Iodide (AV/PI) staining, and graphed as percent of the total cell population. **(D)** Indicated cells were treated with MPA (1 µM, 16 hrs) followed by flow cytometry for phosho-H2AX^S139^ immunostaining, graphed as percent of positive cells. **(E)** Immunoblots from indicated cells treated with MPA (1 µM) for indicated times. **(F, G)** Immunoblots from indicated cells treated for 24 hrs with MPA (1 µM) with or without RMC-5552 (1 nM), rapamycin (20 nM), MK-2206 (2 µM) or SCH-772984 (1 µM). Graphical data are represented as mean ± SEM. (See also Figure S5).

To determine which pathways upstream of mTORC1 confer sensitivity to IMPDH inhibition in these cells, we treated with MPA in the presence or absence of the AKT inhibitor MK-2206 or the ERK1/2 inhibitor SCH-772984. In HSB-2 cells, AKT inhibition resulted in mTORC1 inhibition, and blocked MPA-induced Caspase-3 cleavage, similar to mTORC1 inhibition with RMC-5552 or rapamycin (Figure 5F), suggesting these cells likely have other AKT-activating mutations that confer sensitivity through mTORC1. In contrast, AKT inhibition was not sufficient to block MPA-induced Caspase-3 cleavage in KOPTK-1 cells, and did not inhibit mTORC1 as strongly in these cells (Figure 5G). Instead, Caspase-3 cleavage in these cells was slightly reduced by ERK inhibition, and more strongly reduced by combined inhibition of AKT and ERK, which also robustly inhibited mTORC1 (Figure 5G), suggesting IMPDH inhibitor-sensitivity in these cells is driven by AKT and ERK signaling. These data indicate that mTORC1-activation through AKT and ERK signaling can confer similar sensitivity as PTEN-loss, thus significantly broadening the potential anti-cancer applications for IMPDH inhibitors.

## Discussion

Here we find that PTEN-deficiency in T-ALL confers dependence on the rate-limiting enzyme in guanylate nucleotide synthesis, IMPDH, for cell growth and viability. This dependence stems from uncontrolled AKT/mTORC1 activation, and can be exploited using clinically approved IMPDH inhibitors. Building on previous findings in TSC complex-deficient cells and tumors (28,29), this work demonstrates that mTORC1-activating mutations that lie upstream of the TSC complex can confer robust dependence on IMPDH. mTORC1-activating mutations that lie upstream of the TSC complex are much more prevalent in cancer than mutations in the TSC complex itself (54–56), and include mutations in the PI3K/AKT pathway, the RAS pathway, receptor tyrosine kinases, and others, thus significantly broadening the anti-cancer potential of this metabolic vulnerability.

Chronic mTORC1 activation engages robust negative-feedback loops that block hormone/growth factor signaling to PI3K/AKT. Thus, AKT is inactive in TSC complex-deficient cells (57,58), whereas PTEN-deficiency circumvents this feedback inhibition to activate both AKT and mTORC1. This is a key distinction between TSC complex-deficient and PTEN-deficient cells, given that AKT has effects on metabolism, cell growth, proliferation, and survival that are independent of its effects on mTORC1 (19,20). Our data demonstrate that IMPDH inhibition can be effective in cancer cells that have active AKT, and that AKT contributes to the sensitivity through activation of mTORC1.

CRISPR screening data from the Cancer Dependency Map (DepMap, https://depmap.org/portal/) indicate that leukemia and lymphoma cells are dependent on IMPDH for growth and viability, with T-cell acute lymphoblastic leukemia as the top hit (Supplemental Figure S1A). This is consistent with the well-established immunosuppressive nature of IMPDH inhibitors, which, combined with enhanced IMPDH dependence conferred by mTORC1 activation, raises the possibility that IMPDH inhibitors could potentially be repurposed for these cancers. Future work will determine whether PTEN-deficiency can similarly sensitize solid tumors to IMPDH inhibition, including prostate cancer, endometrial cancer, and gliomas, where PTEN-loss is common (54–56).

Our data in PTEN-wild type HSB-2 and KOPTK-1 cells also suggest that other mTORC1-activating mutations upstream of AKT and ERK could potentially sensitize T-ALL cells to IMPDH inhibition (Figure 5F, 5G). Both AKT and ERK are known to activate mTORC1 by inhibiting the TSC complex, and we demonstrated that sensitivity in these cells is dependent on mTORC1. These results suggest that IMPDH inhibitor efficacy in T-ALL is determined by mTORC1 status, and may not be limited to just PTEN-deficient cells. Given the prevalence of mTORC1 activation in leukemias and lymphomas, including downstream of common oncogenic fusions and mutations in other components of the PI3K/AKT pathway (16,25,59), future work will determine the full translational potential of this metabolic vulnerability in T-ALL. Our findings also raise the possibility that mutations upstream of AKT and ERK could be drivers of IMPDH inhibitor-sensitivity in other cancers as well.

Even among the T-ALL cell lines that have uncontrolled mTORC1 activation, 2 of the lines that are most sensitive to IMPDH inhibition, MOLT-3 and KOPTK-1, show clear p53 induction in response to MPA (Figures 3A, 5E). Induction of apoptosis is also detected earlier in these cells, evident by Caspase-3 cleavage within 18 hrs of MPA treatment, compared to 48 hrs in other PTEN-deficient cells and HSB-2 cells (Figures 3A, 5E). Given the well-established roles of p53 in controlling cell cycle progression, the cellular response to DNA damage, and induction of apoptosis, p53 status could be an additional determinant of sensitivity to IMPDH inhibition, in T-ALL, in addition to PTEN/mTORC1 status. Importantly, p53 loss-of-function is less common in ALLs than in many other cancers (60), suggesting additional stratification based on p53 status could possibly predict exceptional responders.

Although the clinically approved IMPDH inhibitors mizoribine and MPA/MMF have similar immunomodulatory effects in patients, and in mice, their therapeutic efficacy in pre-clinical tumor/cancer models differs substantially for reasons that are not fully understood (29,61–63). Variable mizoribine efficacy can be attributed to differences in the expression level of adenosine kinase (ADK), which is required to convert mizoribine to the active mizoribine-monophosphate to inhibit IMPDH (29,43,44). MPA is inactivated via glucuronidation *in vivo*, and polymorphisms in UDP-glucuronosyltransferases (UGTs) are associated with altered MPA pharmacokinetics in patients (64,65). Relative to mizoribine, MPA is less selective in targeting TSC complex-deficient tumor cells compared to control cells, and has greater effects on adenylate nucleotide levels under treatment conditions that have similar effects on guanylate levels (29). MPA also more readily reduces mTORC1 activity, which could possibly limit its efficacy in exploiting mTORC1-dependent metabolic vulnerabilities (29). Thus, although MPA demonstrates robust efficacy *in vitro*, and MMF can show modest efficacy *in vivo* which is superior to mizoribine when ADK levels are low, mizoribine appears to outperform MPA/MMF when ADK levels are not limiting.

We also evaluated the newly identified IMPDH inhibitor gliocidin (42) in our T-ALL models, which exhibited selective anti-leukemic activity in PTEN-deficient T-ALL cells relative to PTEN-expressing cells (Supplemental Figure S1D), providing rationale for further testing of gliocidin. Future work is also necessary to determine whether IMPDH inhibitors can enhance the efficacy of current therapy regimens, especially when combined with therapies that target other nodes in nucleotide metabolism in leukemia (e.g. methotrexate, mercaptopurine) and other cancers. By further defining mechanisms of sensitivity and resistance, and tailoring specific IMPDH inhibitors to the most appropriate cancer settings, there seems to be strong potential for these safe and well-tolerated therapeutics to benefit stratified patient populations.

## Materials and Methods

### Cell lines

TALL-1 (DSMZ #ACC 521), HPB-ALL (DSMZ #ACC 483), DND-41 (DSMZ #ACC 525), ALL-SIL (DSMZ #ACC 511), RPMI-8402 (DSMZ #ACC 290), LOUCY (DSMZ #ACC 394), JURKAT (DSMZ #ACC 282), CCRF-CEM (DSMZ #ACC 240), KOPTK-1 (Cellosaurus RRID #CVCL_4965), HSB-2 (ATCC #ACC 435), MOLT-3 (ATCC #CRL-1552), and Ba/F3 cells (DMSZ #ACC 300) were grown in RPMI-1640 (Corning #10-040-CV), supplemented with 10% heat-inactivated fetal bovine serum (ThermoFisher Scientific # A5256701) and 1% penicillin-streptomycin (Corning #30-002-Cl). Ba/F3 cells were a kind gift from Dr. Lina Obeid (Stony Brook University), authenticated by short-tandem repeat (STR) profiling at IDEXX BioAnalytics (Columbia, MO), and grown in media supplemented with 10 ng/mL IL-3 (R&D systems #403-ML010). To generate Ba/F3 cells stably expressing non targeting sgRNA (sgCtl) or sgRNA targeting PTEN (sgPTEN), HEK293T cells were transfected with pLentiCRISPR v2 control or sgPTEN (gRNA sequence: GCAGCAATTCACTGTAAAGC, GenScript USA Inc.) along with packaging vectors psPAX2 and pVSVG in a ratio of 4:3:1. Viral supernatant was collected 48 hours post transfection and 5 x 10^5^ Ba/F3 cells were transduced in the presence of 10 μg/ml polybrene. Media was changed 24 hours post transduction. Infected cells were selected with 1 μg/ml puromycin to generate stable cell lines. JURKAT cells with doxycycline-inducible PTEN expression were grown in 1 µg/mL doxycycline (46).

### Mouse studies

All the animal experiments were approved by Rutgers Institutional Animal Care and Use Committee (IACUC). The GEMM model used in Figure 4A-C and Supplemental Figure S4A-D, was described previously (52). Briefly, to induce T-ALL development, bone marrow hematopoietic progenitors from tamoxifen-inducible conditional PTEN-knockout mice (Rosa26^Cre-ERT2/+^PTEN^fl/fl^) were infected with retrovirus expressing a constitutively active oncogenic NOTCH1 receptor mutant (L1601P Δ-PEST), along with a luciferase reporter, and then transplanted into isogenic recipient mice. T-ALL cells from these mice were then injected into irradiated secondary recipients via retro-orbital injection, and treated with vehicle or tamoxifen (5 mg per mouse by i.p. injection) 2 days later to generate PTEN-expressing or PTEN-deleted T-ALL, respectively. Beginning 4 days after tamoxifen injection, mice were treated daily with vehicle, mizoribine (100 mg/kg/day for the first week and then 75 mg/kg/day thereafter, by i.p. injection, resuspended in saline) of MMF (200 mg/kg/day by oral gavage, resuspended in 5% dextrose in saline, pH adjusted to 4-4.5 with HCl) until sacrifice upon reaching humane endpoint. *In vivo* luciferase imagining was performed using the In Vivo Imaging System (IVIS, Lumina LT Series III, PerkinElmer).

For the xenograft studies, 2 million MOLT-3 cells were injected into the tail vein of 6–7-week-old NOD.Cg-Prkdc^scid^ Il2rg^tm1Wjl^ /SzJ mice (Jackson Laboratory #005557). Beginning 6 days later, mice were treated as above with vehicle, mizoribine (75 mg/kg/day by i.p. injection) or MMF (200 mg/kg/day by oral gavage) until sacrifice upon reaching humane endpoint.

### Chemical compounds

The following compounds were added into culture medium at final concentrations indicated in the figure legends: Rapamycin (Sigma #553210) in DMSO, RMC-5552 (MedChemExpress #HY-132168) in DMSO, MK-2206 (Selleckchem #S1078) in DMSO, SCH-772984 (MedChemExpress #HY-50846) in DMSO, mycophenolic acid (Sigma #M3536) in DMSO, mizoribine (Selleckchem #S1384 and Sigma #M3047) in water, gliocidin (MedChemExpress #HY-124778) in DMSO, Q-VD-Oph (Sigma # SML0063) in DMSO, guanosine (Sigma #G6752) in DMSO.

### Cell viability assays

Cell viability was measured using the Cell Titer Glo Luminescent Cell Viability Assay (Promega #G7573) according to manufacturer’s instructions. Annexin V/PI staining was performed using the Dead Cell Apoptosis Kit (ThermoFisher Scientific #V13245), according to manufacturer’s instructions. Staining was measured with a Gallios Flow Cytometer (Beckman Coulter) and analyzed with Kaluza Analysis 2.1 software.

### Metabolite analysis by LC-MS/MS

Equal number of T-ALL cells were centrifuged to pellet (1000 rpm, 5 minutes, RT), media was aspirated, and metabolites were extracted in 1 mL ice cold extraction buffer (40% methanol: 40% acetonitrile: 20% water with 0.1 M formic acid) per 4 million cells, for 5 minutes on ice, followed by quenching with ammonium carbonate (15% w/v in HPLC grade water) as previously described (66,67). Metabolite extracts were centrifuged (15000xg, 10 minutes, 4°C) and supernatant collected and stored at -80°C before LC-MS/MS analysis. LC-MS/MS was performed on a Q Exactive PLUS hybrid quadrupole–Orbitrap mass spectrometer (ThermoFisher Scientific) coupled to a hydrophilic interaction chromatography (HILIC) platform. Chromatographic separation was carried out on a Vanquish Horizon UHPLC system using an XBridge BEH Amide column (150 mm × 2.1 mm, 2.5 μm; Waters, Milford MA) equipped with the corresponding XP VanGuard pre-column. The LC gradient consisted of solvent A (95:5 H₂O: acetonitrile containing 20 mM ammonium acetate and 20 mM ammonium hydroxide, pH 9.4) and solvent B (20:80 H₂O: acetonitrile with the same buffer composition as A). The gradient program was as follows: 0 min, 100% B; 3 min, 100% B; 3.2 min, 90% B; 6.2 min, 90% B; 6.5 min, 80% B; 10.5 min, 80% B; 10.7 min, 70% B; 13.5 min, 70% B; 13.7 min, 45% B; 16 min, 45% B; and 16.5 min, 100% B. The flow rate was maintained at 300 μL/min, with a 5 μL injection volume and a column temperature of 25°C. Mass spectrometry was performed in negative ion mode with a resolution of 70,000 at m/z 200. The automatic gain control (AGC) target was set to 3 × 10⁶, and full-scan spectra were acquired over an m/z range of 75-1,000. Metabolite features were extracted using MAVEN (68), specifying the appropriate labeled isotopes and applying a mass-accuracy window of 5 ppm.

### Immunoblotting and antibodies

Cells were lysed in triton X-100 lysis buffer (20 mM Tris pH 7.5, 140 mM NaCl, 1 mM EDTA, 10% glycerol, 1% triton X-100, 1 mM DTT, 50 mM NaF) or RIPA buffer (150 mM NaCl, 1% IGEPAL, 0.5% sodium deoxycholate, 0.1%SDS, 50 mM Tris pH8, 1 mM DTT), supplemented with protease inhibitor cocktail (Sigma #P8340), phosphate inhibitor cocktail #2 (Sigma #P5726) and #3 (Sigma #P0044) at 1:100 each. Western blots were performed using the following antibodies at 1:1000 dilution unless otherwise noted: PTEN (CST #9188, 1:2500), Cleaved Caspase-3 (CST #9664), pH2AXS^139^ (CST #9718), CHK1 (CST #2360, 1:2000), pCHK1^S345^ (CST #2348, 1:2000), CHK2 (CST #2662, 1:1500), pCHK2^T68^ (CST #2661), p53 (CST #9282), p21 (CST #2947T), AKT (CST #4691, 1:2000), pAKT^T308^ (CST #9275, #13038), pAKT^S473^ (CST #4060, 1:5000), pGSK-3^S9/21^ (CST #9331), S6K (CST #2708, 1:2000), pS6K^T389^ (CST #9234), GAPDH (CST #5174, 1:10000), 4E-BP1 (CST #9644, 1:5000), PARP (CST #9542), RPA (CST #52448), pRPA^T21^(Abcam #ab109394, 1:2000), Cyclin E (CST #20808), ERK 1/2 (CST #9102, 1:1000), pERK^T202/Y204^ (CST #9106, 1:1500).

### RNA synthesis measurements

RNA synthesis rate was quantified using the global RNA Synthesis Assay Kit (Abcam #ab228561) according to manufacturer’s instructions. Cells were cultured as above and treated with or without Actinomycin D for 3 hours followed by incubation with 5-ethynyl-uridine (5-EU) for 1 hour to label newly synthesized RNA. Cells were then fixed, permeabilized, and the fluorescent azide dye was conjugated as per manufacturer’s instructions. Staining was analyzed on a Gallios Flow Cytometer (Beckman Coulter) and analyzed using Kaluza Analysis 2.1 software.

### Cell cycle profiling

Cells were spun to pellet (1000 rpm, 5 minutes, RT) followed by washing in 1X PBS + 0.1% BSA (Sigma #A7030). Cells were then fixed in 100% ethanol (pre-chilled to -20°C) for 15 minutes on ice, followed by washing with 1X PBS. Cells were then stained with propidium iodide-RNase solution (CST #4087) for 1 hour at room temperature. PI staining intensity was measured on a Gallios Flow Cytometer (Beckman Coulter) and data were analyzed using Kaluza Analysis 2.1 software.

### phospho-H2AX immunostaining

Cells were spun to pellet (1000 rpm, 5 minutes, RT), washed with PBS + 0.1% BSA (Sigma #A7030) and spun again. Cells were then fixed in 4% formaldehyde (ThermoFisher Scientific #28908) solution for 15 minutes at RT and washed in PBS + 0.1% BSA. The cells were then permeabilized in 100% methanol (pre-chilled to -20°C) for 10 minutes on ice, washed as above, and incubated with phospho-H2AX^S139^ primary antibody (CST #9718, 1:50 in 1X PBS + 0.1% BSA) for 1 hr at RT. Cells were then washed as above, incubated with Cy3-conjugated secondary antibody (Jackson ImmunoResearch #111-165-144, 1:1000) for 30 minutes at room temperature, washed again, spun to pellet, counterstained with 0.5 μg/mL DAPI (ThermoFisher #D1306) for 1 minute, and washed again. Fluorescence intensity was measured on a Gallios Flow Cytometer (Beckman Coulter), and data were analyzed using Kaluza Analysis 2.1 software.

### Quantification and Statistical Analysis

Graphical data are represented as mean of at least three biological replicates ±SEM, unless otherwise indicated in the figure legends. Graphs were plotted and analyzed in Graph Pad Prism 10. p-vales for pairwise comparisons between two groups were determined using an unpaired Welch’s t-test. p values for comparisons between more than two groups were determined using two-way ANOVA. Statistical details for individual experiments can be found in their respective figure legends.

## Supporting information

Supplementary figures with legends

## Data availability

The data analyzed in Supplementary Figures S1A and S1B were obtained from The Cancer Dependency Map at https://depmap.org/portal/. Metabolomics data were generated at the Rutgers Cancer Institute Metabolomics Core Facility. Derived metabolomics data and other data generated in this study are available from the corresponding author upon request.

## Acknowledgements

We thank Masuda Akther, Nanda Singh, Aruna Jangam, Jay Joshi, Alexandra Grumet, and Ariel Lerner for helpful discussions and technical assistance. We also thank EOHSI (Environment and Occupational Health Sciences Institute) at Rutgers Flow Cytometry Core members Rita Hahn, Jessica Cervelli and Raymond Rancourt for technical assistance. This work was supported by grants from the Ludwig Institute for Cancer Research -Princeton Branch (to A.J.V.), the Leukemia Research Foundation (New Investigator Research Grant to A.J.V.), the Pediatric Cancer and Blood Disorders Research Center at the Rutgers Cancer Institute (Research Grant to A.J.V.), and US National Institutes of Health (R01CA232246 to W.X.Z.). Work in the laboratory of D.H. is supported by the US National Institutes of Health (R01CA236936 and R01CA285513) and Blood Cancer United, formerly The Leukemia & Lymphoma Society (Scholar Award 1386-23). The Rutgers Cancer Institute metabolomics core facility is supported by the US National Institutes of Health (NCI-CCSG P30CA072720-6852).

## Notes

Conflict of Interest Statement: The authors declare no potential conflicts of interest.

### Competing Interest Statement

The authors have declared no competing interest.

