## Supplementary figures with legends for "PTEN-loss confers dependence on the guanylate synthesis enzyme IMPDH in T-cell acute lymphoblastic leukemia"

Fig S1:

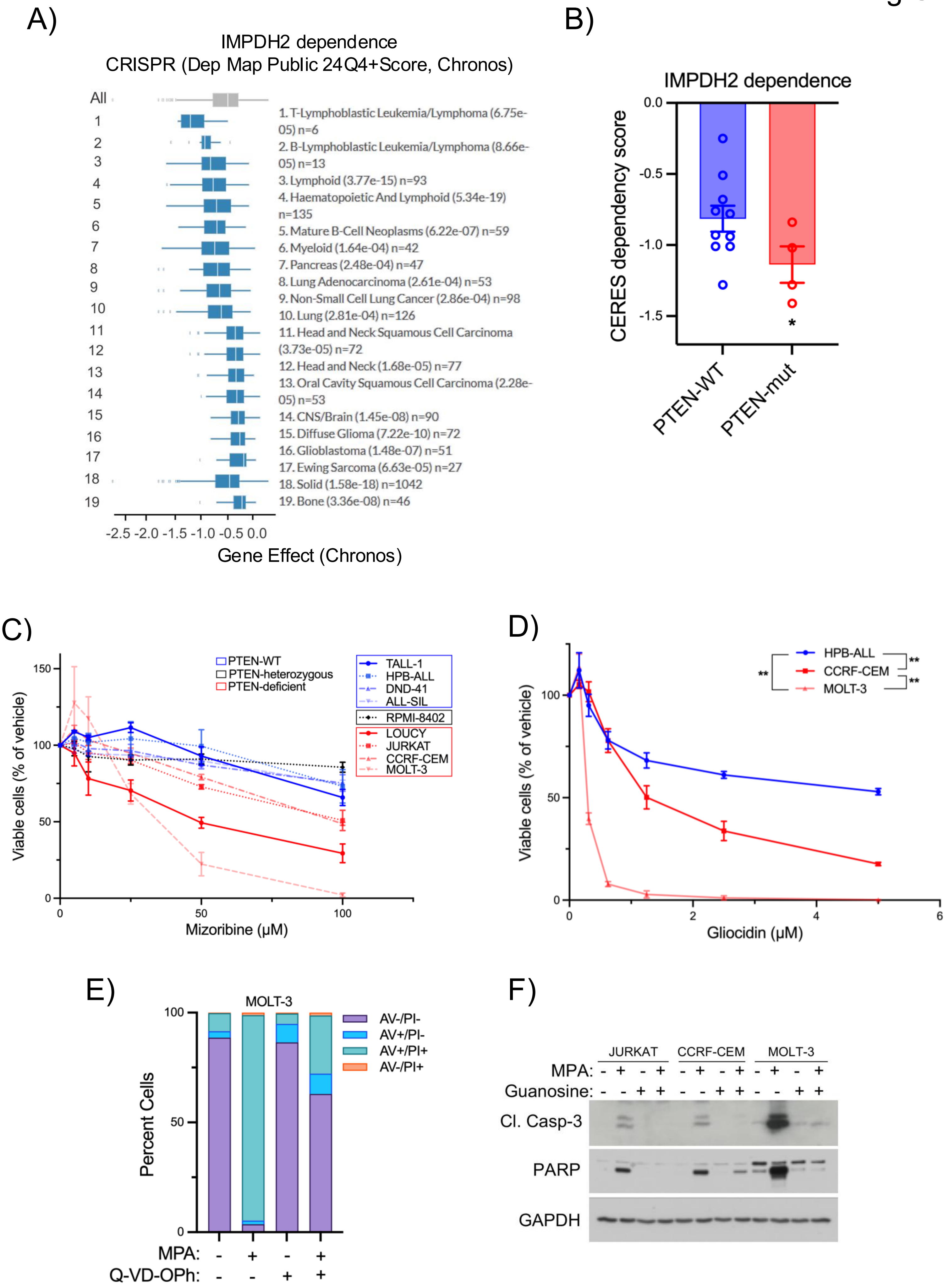

A)
IMPDH2 dependence
CRISPR (Dep Map Public 24Q4+Score, Chronos)
All
1. T-Lymphoblastic Leukemia/Lymphoma (6.75e-05) n=6
2. B-Lymphoblastic Leukemia/Lymphoma (8.66e-05) n=13
3. Lymphoid (3.77e-15) n=93
4. Haematopoietic And Lymphoid (5.34e-19) n=135
5. Mature B-Cell Neoplasms (6.22e-07) n=59
6. Myeloid (1.64e-04) n=42
7. Pancreas (2.48e-04) n=47
8. Lung Adenocarcinoma (2.61e-04) n=53
9. Non-Small Cell Lung Cancer (2.86e-04) n=98
10. Lung (2.81e-04) n=126
11. Head and Neck Squamous Cell Carcinoma (3.73e-05) n=72
12. Head and Neck (1.68e-05) n=77
13. Oral Cavity Squamous Cell Carcinoma (2.28e-05) n=53
14. CNS/Brain (1.45e-08) n=90
15. Diffuse Glioma (7.22e-10) n=72
16. Glioblastoma (1.48e-07) n=51
17. Ewing Sarcoma (6.63e-05) n=27
18. Solid (1.58e-18) n=1042
19. Bone (3.36e-08) n=46
-2.5 -2.0 -1.5 -1.0 -0.5 0.0
Gene Effect (Chronos)
B)
IMPDH2 dependence
CERES dependency score
0.0
-0.5
-1.0
-1.5
*
PTEN-WT
PTEN-mut
C)
PTEN-WT
PTEN-heterozygous
PTEN-deficient
TALL-1
HPB-ALL
DND-41
ALL-SIL
RPMI-8402
LOUCY
JURKAT
CCRF-CEM
MOLT-3
Viable cells (% of vehicle)
150
100
50
0
Mizoribine (µM)
0
50
100
D)
HPB-ALL
CCRF-CEM
MOLT-3
**
**
**
Viable cells (% of vehicle)
100
50
0
Gliocidin (µM)
0
2
4
6
E)
MOLT-3
AV-/PI-
AV+/PI-
AV+/PI+
AV-/PI+
Percent Cells
100
50
0
MPA: - + - +
Q-VD-OPh: - - + +
F)
JURKAT
CCRF-CEM
MOLT-3
MPA: - + - + - + - + - + - +
Guanosine: - - + + - - + + - - + +
Cl. Casp-3
PARP
GAPDH

**Figure S1, related to Figure 1:**

**(A, B)** Cancer cell dependence on IMPDH2 from The Cancer Dependency Map (DepMap). **(A)** Gene Effect score for IMPDH2 dependence in cell lines from indicated cancer lineages. Lower scores indicate more dependence on IMPDH2. **(B)** CERES dependency score from acute lymphoblastic leukemia cell lines, graphed according to *PTEN* status.

**(C, D)** Indicated T-ALL cells were treated with indicated concentrations of the IMPDH inhibitor mizoribine **(C)**, or gliocidin **(D)** for 72 hrs. Viable cells were measured using the Cell Titer Glo Cell Viability Assay (Promega) and graphed as percent of vehicle-treated cells.

**(E)** MOLT-3 cells (PTEN-deficient) were treated for 72 hours with mycophenolic acid (MPA, 2.5 μM), with or without the pan-Caspase inhibitor Q-VD-Oph (20 μM). Cell death was quantified by Annexin V/ Propidium Iodide (AV/PI) staining and graphed as percent of the total cell population.

**(F)** Immunoblots from indicated T-ALL cells treated with vehicle or MPA (1 μM, 48 hrs)**,** with or without supplementation with exogenous guanosine (25 μM).

Graphical data are represented as mean ± SEM. *p< 0.05, **p < 0.0001 by Welch's t-test and two-way ANOVA.

Fig S2:

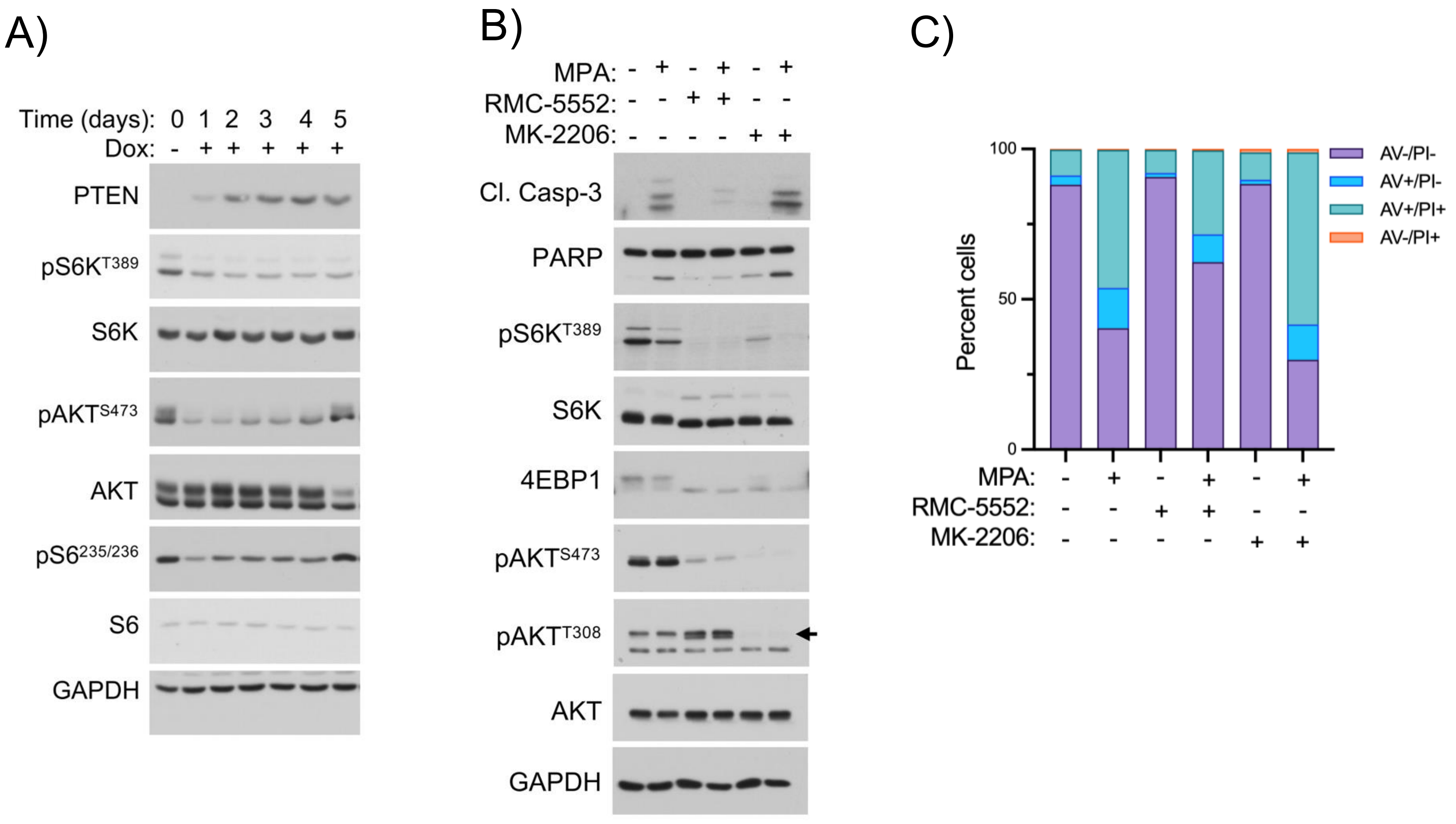
A)
Time (days): 0 1 2 3 4 5
Dox: - + + + + +
PTEN
pS6K T389
S6K
pAKT S473
AKT
pS6 235/236
S6
GAPDH
B)
MPA: - + - + - +
RMC-5552: - - + + - -
MK-2206: - - - - + +
Cl. Casp-3
PARP
pS6K T389
S6K
4EBP1
pAKT S473
pAKT T308
AKT
GAPDH
C)
Percent cells
100
50
0
AV-/PI-
AV+/PI-
AV+/PI+
AV-/PI+
MPA: - + - + - +
RMC-5552: - - + + - -
MK-2206: - - - - + +

**Figure S2, related to Figure 2:**

**(A)** Immunoblots from JURKAT cells with doxycycline-inducible PTEN-addback, treated with doxycycline (Dox, 1 µg/mL) for indicated times.

**(B)** Immunoblots from JURKAT cells treated for 48 hrs with MPA (1 µM) with or without RMC-5552 (1 nM), or MK-2206 (2µM).

**(C)** Jurkat cells were treated for 72 hrs with MPA (2.5 µM) with or without RMC-5552 (1 nM) or MK-2206 (2 µM). Cell death was quantified by Annexin V/ Propidium Iodide (AV/PI) staining and graphed as percent of the total cell population.

Fig S3:

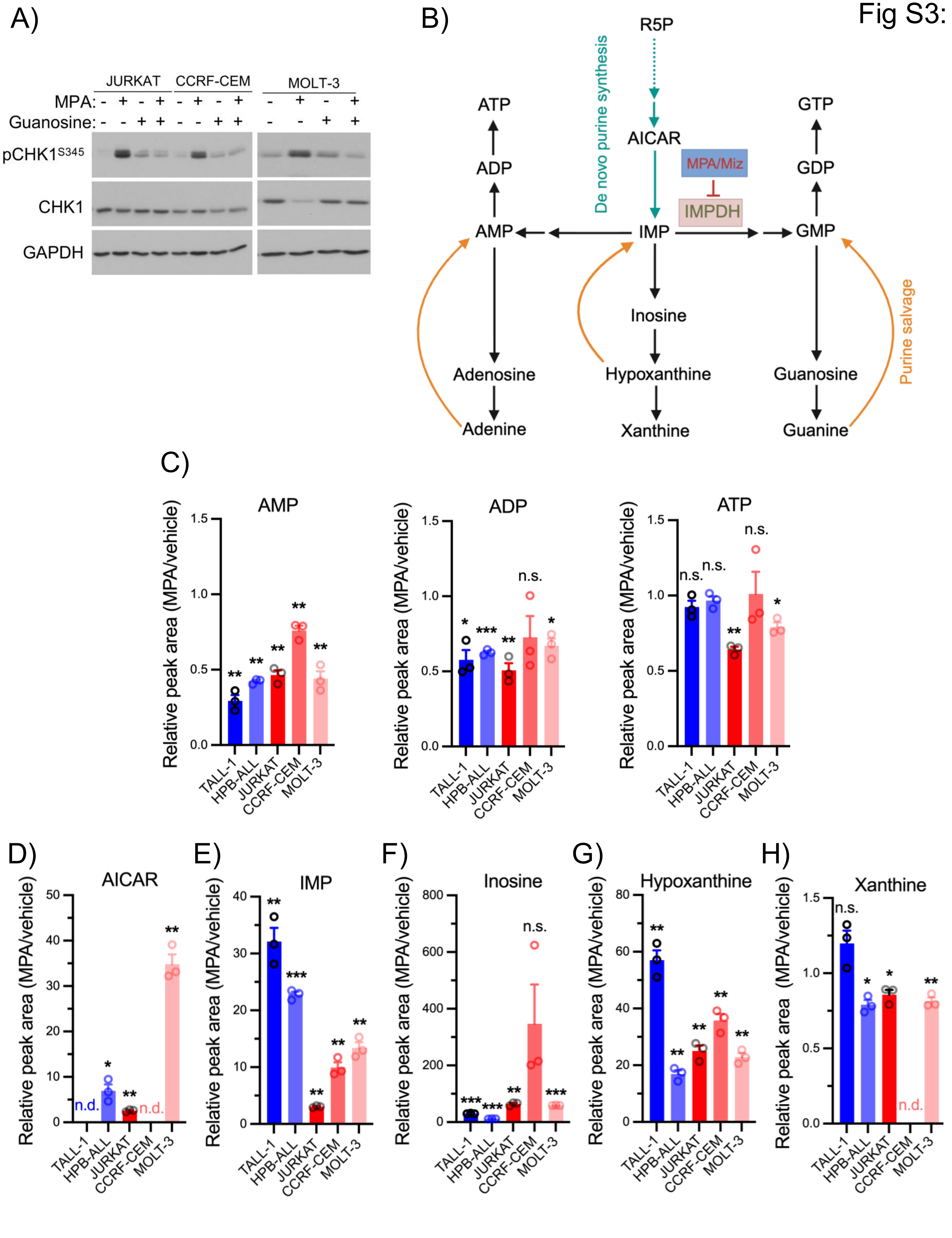
A)
JURKAT
CCRF-CEM
MOLT-3
MPA: - + - + - + - + - + - +
Guanosine: - - + + - - + + - - + +
pCHK1S345
CHK1
GAPDH
B)
R5P
De novo purine synthesis
AICAR
MPA/Miz
IMPDH
ATP
ADP
AMP
IMP
GMP
GDP
GTP
Inosine
Adenosine
Hypoxanthine
Guanosine
Adenine
Xanthine
Guanine
Purine salvage
C)
AMP
ADP
ATP
Relative peak area (MPA/vehicle)
n.s.
TALL-1
HPB-ALL
JURKAT
CCRF-CEM
MOLT-3
D)
AICAR
n.d.
E)
IMP
F)
Inosine
G)
Hypoxanthine
H)
Xanthine

**Figure S3, related to Figure 3:**

**(A)** Immunoblots from indicated cells treated for 24 hrs with MPA (1 µM) with or without supplementation with exogenous guanosine (25 µM).

**(B)** Schematic of *de novo* purine synthesis and salvage pathways.

**(C-H)** Relative abundance of the indicated metabolites, measured by LC-MS/MS, in cells treated with MPA (1 µM, 6 hours), graphed normalized to vehicle-treated cells.

Graphical data are represented as mean ± SEM. *$p < 0.05$, **$p < 0.005$, ***$p < 0.0005$ by Welch's t-test.

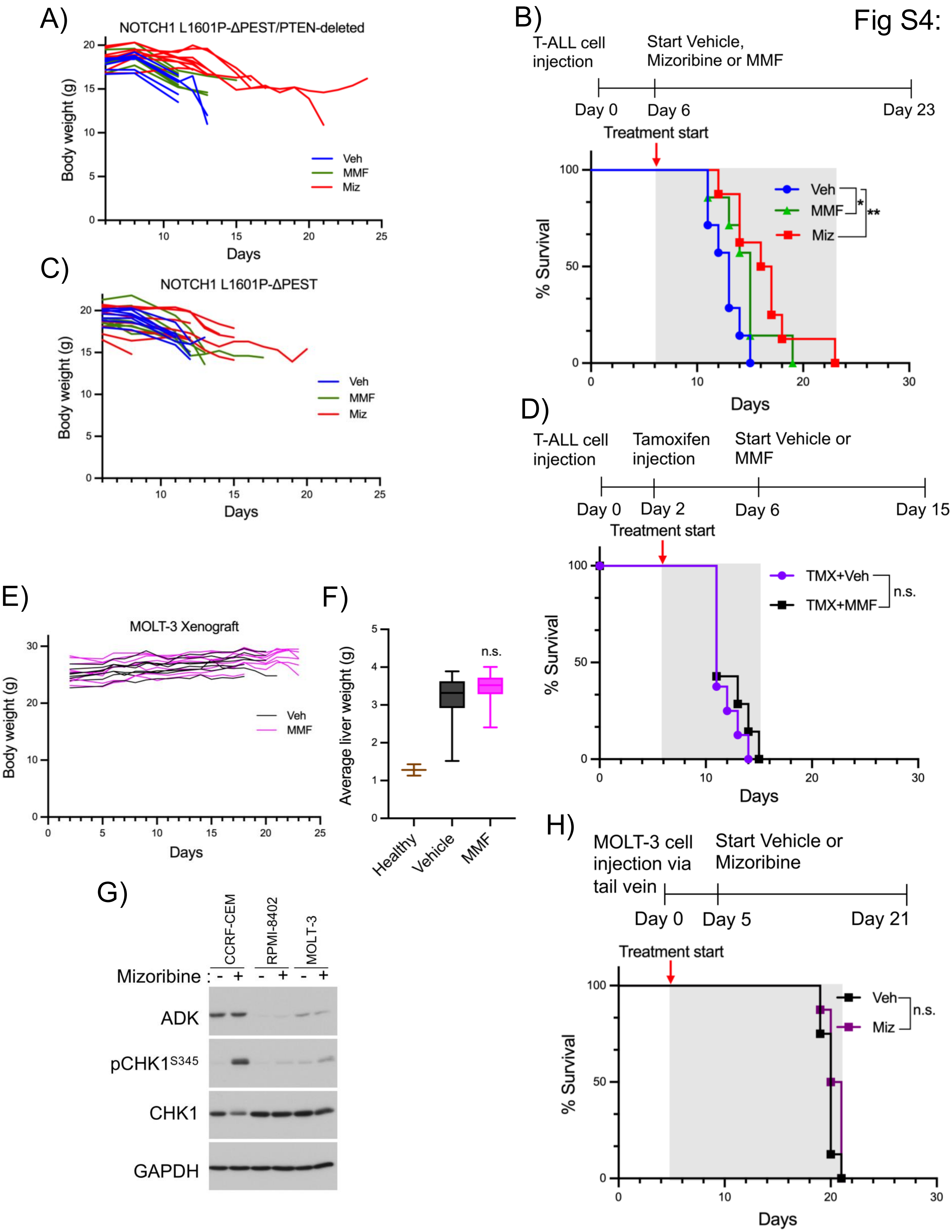

Fig S4:
A)
NOTCH1 L1601P-ΔPEST/PTEN-deleted
Body weight (g)
Days
Veh
MMF
Miz
B)
T-ALL cell injection
Start Vehicle, Mizoribine or MMF
Day 0
Day 6
Day 23
Treatment start
% Survival
Days
Veh
MMF
Miz
*
**
C)
NOTCH1 L1601P-ΔPEST
Body weight (g)
Days
Veh
MMF
Miz
D)
T-ALL cell injection
Tamoxifen injection
Start Vehicle or MMF
Day 0
Day 2
Day 6
Day 15
Treatment start
% Survival
Days
TMX+Veh
TMX+MMF
n.s.
E)
MOLT-3 Xenograft
Body weight (g)
Days
Veh
MMF
F)
Average liver weight (g)
n.s.
Healthy
Vehicle
MMF
G)
CCRF-CEM
RPMI-8402
MOLT-3
Mizoribine :
ADK
pCHK1S345
CHK1
GAPDH
H)
MOLT-3 cell injection via tail vein
Start Vehicle or Mizoribine
Day 0
Day 5
Day 21
Treatment start
% Survival
Days
Veh
Miz
n.s.

**Figure S4, related to Figure 4:**

**(A-D)** To induce T-ALL development, bone marrow hematopoietic progenitors from tamoxifen-inducible conditional PTEN-knockout mice ($Rosa26^{Cre\text{-}ERT2/+}PTEN^{fl/fl}$) were infected with retrovirus expressing a constitutively active oncogenic NOTCH1 receptor mutant (L1601P Δ-PEST), along with a luciferase reporter, and then transplanted into isogenic recipient mice. T-ALL cells from these mice were then injected into irradiated secondary recipients (via retro-orbital injection)**.** 2 days later, mice were treated with vehicle or tamoxifen to generate PTEN-expressing or PTEN-deleted T-ALL and treated daily with vehicle, mycophenolate mofetil (MMF, 200 mg/kg/day), or mizoribine (100 mg/kg/day for the first week and then 75 mg/kg/day thereafter) beginning 4 days after vehicle or tamoxifen treatment. n=8 mice per group. **(A)** Body weights of mice from Figure 4A. Each line represents an individual mouse. **(B)** Kaplan-Meier survival plot from mice bearing PTEN-expressing T-ALL treated with vehicle, MMF, or mizoribine. **(C)** Body weights of mice from Figure S4B. Each line represents an individual mouse. **(D)** Kaplan-Meier survival plot from mice bearing PTEN-deleted T-ALL treated with vehicle or mizoribine.

**(E)**. Body weights of mice from Figure 4D. Each line represents an individual mouse.

**(F)** Average liver weights from mice in Figure 4D compared to healthy non-leukemic controls.

**(G)** Immunoblots on indicated T-ALL cells treated with vehicle or mizoribine (250 μM, 24 hrs).

**(H)** T-ALL xenografts were established via tail vein injection of MOLT-3 cells. Beginning 5 days later, mice were treated daily with vehicle or mizoribine (75 mg/kg/day) until succumbing. Kaplan-Meier survival plots are shown. n=8 mice per group.

Fig S5:

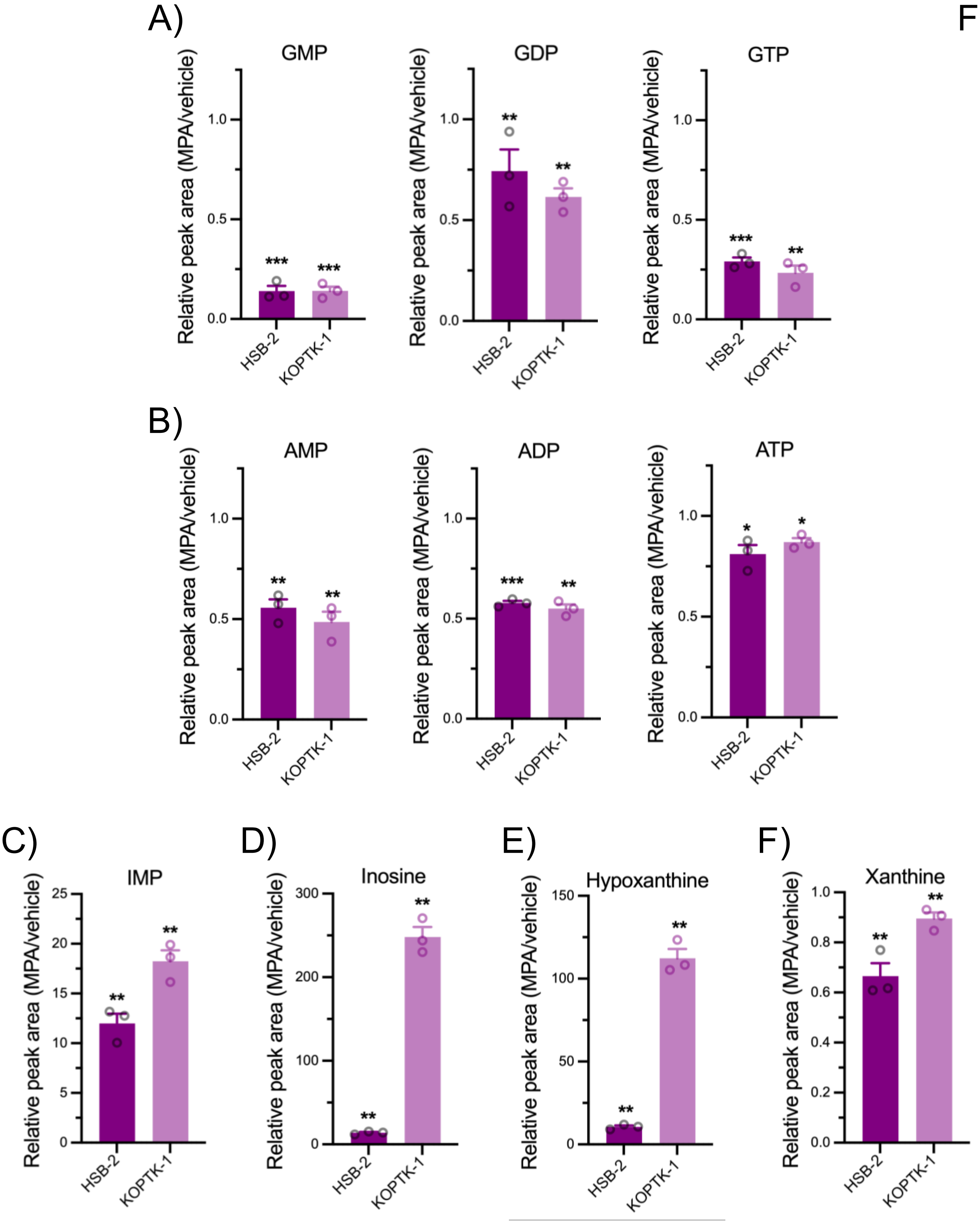
A)
GMP
GDP
GTP
B)
AMP
ADP
ATP
C)
IMP
D)
Inosine
E)
Hypoxanthine
F)
Xanthine
Relative peak area (MPA/vehicle)
HSB-2
KOPTK-1

**Figure S5, related to Figure 5:**

**(A-F)** Relative abundance of the indicated metabolites, measured by LC-MS/MS, in cells treated with MPA (1 μM, 6 hours), graphed normalized to vehicle-treated cells.

Graphical data are represented as mean ± SEM.*$p < 0.05$, **$p < 0.005$, ***$p < 0.0005$ by Welch's t-test.
